# The cytoplasmic DNA sensor cGAS promotes mitotic cell death

**DOI:** 10.1101/168070

**Authors:** Christian Zierhut, Hironori Funabiki

**Affiliations:** Laboratory of Chromosome and Cell Biology, The Rockefeller University, New York, New York, 10065, USA.

## Abstract

The cyclic GMP-AMP (cGAMP) synthase cGAS counteracts infections by detecting and binding foreign cytoplasmic DNA^1^. DNA-induced synthesis of cGAMP activates innate immune signalling and apoptosis through the cGAMP receptor STING and the downstream effector IRF3^1–7^. During interphase the nuclear envelope protects chromosomal self-DNA from cGAS, but the consequences of exposing chromosomes to cGAS following mitotic nuclear envelope disassembly are unknown. Here we demonstrate that cGAS associates with chromosomes during mitosis and binds nucleosomes with even higher affinity than naked DNA in vitro. Nucleosomes nevertheless competitively inhibit the DNA-dependent stimulation of cGAS, and accordingly, chromosomal cGAS does not affect mitotic progression under normal conditions. This suggests that nucleosomes prevent the inappropriate activation of cGAS during mitosis by acting as a signature of self-DNA. During prolonged mitotic arrest, however, cGAS becomes activated to promote cell death, limiting the fraction of cells that can survive and escape mitotic arrest induced by the chemotherapeutic drug taxol. Induction of mitotic cell death involves cGAMP synthesis by cGAS, as well as signal transduction to IRF3 by STING. We thus propose that cGAS plays a previously unappreciated role in guarding against mitotic errors, promoting cell death during prolonged mitotic arrest. Our data also indicate that the cGAS pathway, whose activity differs widely among cell lines, impacts cell fate determination upon treatment with taxol and other anti-mitotic drugs. Thus, we propose the innate immune system may be harnessed to selectively target cells with mitotic abnormalities.

## Main text

Viral or bacterial infections frequently result in the exposure of pathogen DNA to the cytoplasm of infected cells. This DNA can activate the innate immune system, which induces inflammation, and purges infected cells^1^. An important cytoplasmic DNA sensor is cGAS^4^, which, once bound to DNA, synthesises a cyclic GTP-ATP dinucleotide, cGAMP^1,4,5,8,9^. cGAMP binds to and activates the adapter STING^1,5,10^, ultimately leading to inflammation by triggering the activation of the transcription factors IRF3 and NFκB^1,4,5^. The cGAS pathway can also induce apoptosis, mediated by transcriptional and non-transcriptional mechanisms^2,3,6,7^. The discovery of this cytoplasmic DNA sensor raises the question of how cells ensure that self-DNA does not normally trigger cGAS activation. During interphase, chromosomal self-DNA is protected by the nuclear envelope^11,12^ but it is unknown how cGAS responds during mitosis, when the nuclear envelope disassembles.

We previously hypothesised that nucleosomes, which are uniquely formed on eukaryotic genomic DNA, could be used to distinguish self-DNA from pathogenic DNA^13,14^. Therefore, we tested if cGAS could interact with nucleosomal DNA. Unexpectedly, pull-down assays of^35^ S-methionine-labelled cGAS showed that cGAS bound *in vitro* assembled nucleosomes^13^ much better than naked DNA (Fig. 1a). Even in M-phase *Xenopus* egg extracts, where nucleosome-dependent processes can be assessed in a physiological context^13^, nucleosomes did not inhibit cGAS binding (Extended Data Fig. 1). Gel-shift assays of recombinant cGAS (Extended Data Fig. 2) further demonstrated that cGAS has ~2-fold higher affinity for mononucleosomes than for naked DNA (Fig. 1b, c). Mutations in the cGAS DNA-binding interface^15^ (K173E R176E K407E K411A) greatly reduced affinity for naked DNA, while affecting nucleosome interaction only modestly (Fig. 1d, e), suggesting that mechanisms of cGAS binding to naked DNA and to nucleosomes are distinct.

**Figure 1.**
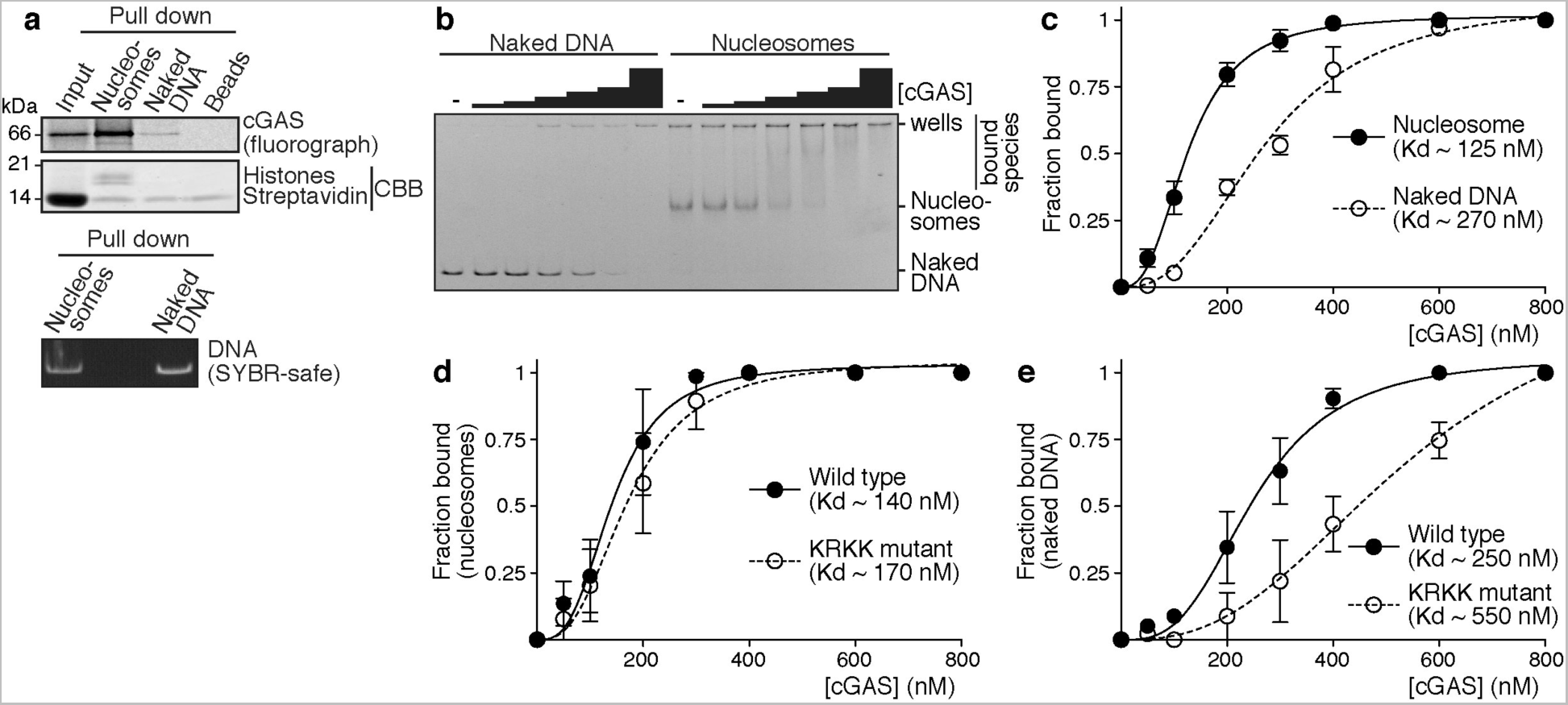
Nucleosomes bind cGAS with higher affinity than naked DNA. **a,** The indicated beads were incubated in buffer containing in vitro generated cGAS. Beads were recovered, washed, and associated proteins were detected by gel electrophoresis and fluorometry. CBB, Coomassie brilliant blue. **b and c,** Analysis of cGAS binding affinity for naked DNA or mononucleosomes by gel mobility shift. **b,** example gel. **c,** quantification. **d and e,** Analysis of the binding affinity of wild type cGAS or cGAS mutated in the DNA binding domain (KRKK mutant) for mononucleosomes (d) or naked DNA (e). All graphs represent mean values and SEM from at least three independent experiments.

Despite the enhanced affinity for cGAS, we found that nucleosomes suppress the cGAMP synthase activity of cGAS. When compared to DNA-bound cGAS, nucleosome-bound cGAS showed a ~3-fold reduction in the apparent catalysis rate (Kcat), as determined by thin-layer chromatography of reactions under saturating conditions of mononucleosomes or naked DNA (Fig. 2a-c). Consistent with the idea that nucleosomes could competitively suppress the capacity of naked DNA to activate cGAS, nucleosomes added to standard reactions containing saturating amounts of naked DNA decreased the apparent Kcat in a manner proportional to the concentration of nucleosomes (Fig. 2d). Altogether, these results suggest that, on chromosomes, nucleosomes could competitively interfere with cGAS activation by nucleosome-free regions.

**Figure 2.**
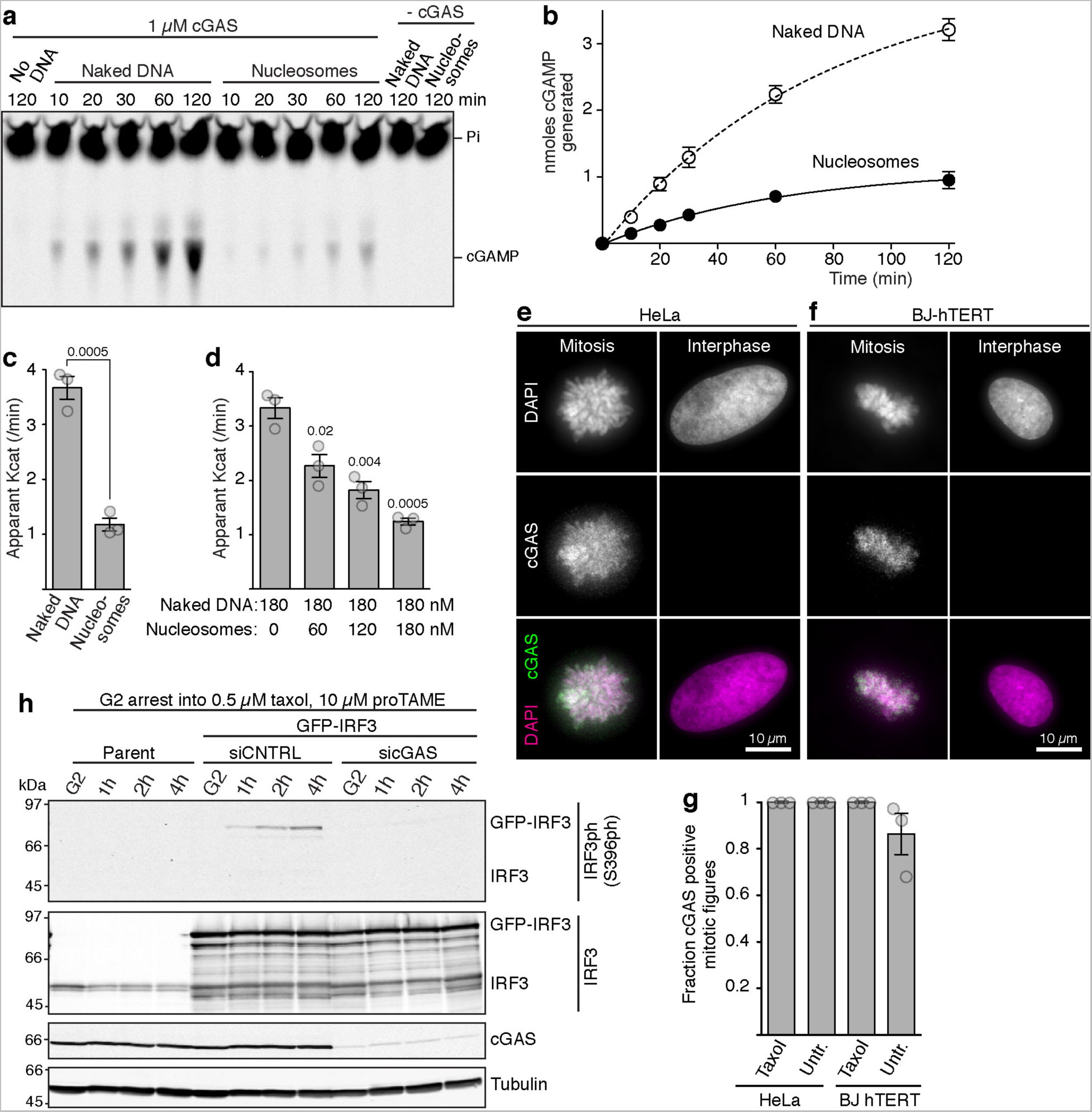
cGAS associates with mitotic chromosomes but nucleosomes limit cGAS activity. **a - b,** Analysis of cGAS activity with naked DNA or mononucleosomes. **a,** Typical example of a cGAS activity assay using either naked DNA or nucleosomes as cGAS stimulator. **b,** Quantification of reactions used to determine Kcat. Mean values and SEM from at least three independent experiments are shown. **c,** Apparent Kcat of cGAS determined with either naked DNA or mononucleosomes. Mean values and SEM from at least three independent experiments are shown. **d,** Apparent Kcat of cGAS in reactions containing naked DNA and increasing concentrations of mononucleosomes. Mean values and SEM from at least three independent experiments are shown. **e - g,** Cells were subjected to immunofluorescence analysis with anti-cGAS antibodies. **e and f,** Example images of cells in unperturbed mitosis. **g,** Quantification of the fraction of mitotic figures that are positive for cGAS localization on chromosomes in either untreated cells (Untr.) or cells treated with 500 nM taxol for 4 h. Shown are mean values with SEM from three experiments (n = 50 each). **h,** Western blot analysis of IRF3 phosphorylation in HeLa cells of the indicated type harvested either in G2 or after the indicated times during arrest in 500 nM taxol, 10 µM proTAME. siCNTRL, control siRNA.

Given our findings that cGAS binds nucleosomes, we speculated that cGAS can freely associate with chromosomes when they are exposed to the cytoplasm. Indeed, similar to a recent report in mouse cells^16^, immunofluorescence analysis demonstrated that cGAS associates with mitotic chromosomes in HeLa cells and in telomerase-immortalized human BJ hTERT fibroblasts (Fig. 2e-g), both during unperturbed mitosis, and during mitotic arrest induced with the chemotherapeutic microtubule-stabilizing drug taxol (paclitaxel). This signal was specific, as it was not observed after siRNA-mediated knockdown of cGAS (Extended Data Fig. 3a, b). Live-imaging of GFP-tagged cGAS (GFP-cGAS) in HeLa Flp-In T-REx cells^17,18^, expressing inducible GFP-cGAS from a single genomic locus, showed that cGAS associates with chromosomes immediately after nuclear envelope break down (Extended Data Fig. 3c). In contrast, cGAS was largely absent from interphase nuclei, with the exception of a subset of micronuclei (Extended Data Fig. 3d, e), structures whose nuclear envelopes are prone to rupture^19,20^.

To determine whether chromosome-bound cGAS could activate the STING pathway during mitosis, we monitored mitotic IRF3 phosphorylation at Ser396 (IRF3ph), a widely used indicator of cGAS activation, in cells arrested in mitosis using taxol. In agreement with the limited cGAS activity on nucleosomes, we found little evidence for IRF3ph on the endogenous protein (Fig. 2h), but mild overexpression of GFP-tagged IRF3 revealed a slow cGAS-dependent increase in IRF3ph during mitotic arrest, but not in interphase (Fig. 2h).

As nucleosomes limit DNA-dependent cGAS activity, cGAS binding to chromosomes may not have consequences during normal mitosis, which completes within an hour, whereas cGAS signalling occurs over multiple hours^4^ (Fig. 2h). Indeed, we found that cGAS depletion neither affected viability or duration of unperturbed mitosis (Extended Data Fig. 4a, b). However, when cells are arrested in mitosis, even low cGAS activity may have functional consequences. Cells arrested in mitosis stochastically undergo one of two fates, death in mitosis or slippage into interphase. The timing of these events differs substantially between cells, and cell fate profiles also vary widely between cell lines^21^. The decision between death and slippage may carry particular weight when tumour cells undergo anti-mitotic chemotherapy, such as taxol. However, the mechanisms underlying the variation in this fate decision are poorly understood^22,23^. Because cGAS protein levels vary drastically among cell lines (Extended Data Fig. 5a), and the cGAS pathway may induce apoptosis independently of transcriptional induction^2,3,7,24^, we tested whether cGAS affects the decision between slippage and death.

We first treated seven breast cancer cell lines (Extended Data Fig. 5a) with a clinically relevant taxol concentration (10 nM)^25,26^. All three cell lines (CAL51, T47D, BT474) that did not express cGAS were able to survive mitotic arrest under this condition, whereas three out of four cell lines expressing cGAS (HCC1143, MDA-MB-157, BT549) were prone to die in mitosis (Extended Data Fig. 5b). Similar effects were observed with a high taxol dose (0.5 µM; Extended Data Fig. 5c). No difference in mitotic viability was observed in the absence of taxol, suggesting that the observed cell death depends on mitotic arrest (Extended Data Fig. 5d). siRNA-mediated depletion of cGAS reduced mitotic mortality by ~50% in the three most sensitive cell lines (Extended Data Fig. 5e, f), further suggesting an involvement of cGAS in the decision between death and slippage in these breast cancer cells.

To characterise the mechanism by which cGAS promotes mitotic death in detail, we focussed on two cell lines, HeLa Flp-In T-REx cell lines, which were used to inducibly express siRNA-resistant, wild type, or mutant GFP-cGAS genes, and untransformed BJ-hTERT fibroblasts, which have been extensively used to analyse the cGAS-STING pathway^8,27^. Similar to our observations in breast cancer cells, cGAS depletion resulted in a reduction in mitotic cell death in HeLa cells treated with 10 nM taxol (Fig. 3a, b). This effect could be a consequence of either enhancing the rate of slippage, or of a delay in the advent of cell death, thus giving cells more time to slip. To distinguish between these possibilities, we monitored the impact of cGAS depletion on the timing of cell death during taxol treatment. Because a change in slippage probability indirectly affects the apparent timing of mitotic cell death^28^, slippage events were blocked by using a higher concentration of taxol (500 nM) in conjunction with the anaphase-promoting complex inhibitor proTAME^29^. This setup, combined with time-lapse microscopy, allowed us to determine individual mitotic lifespans, solely dictated by the timing of death, but not by slippage.

**Figure 3.**
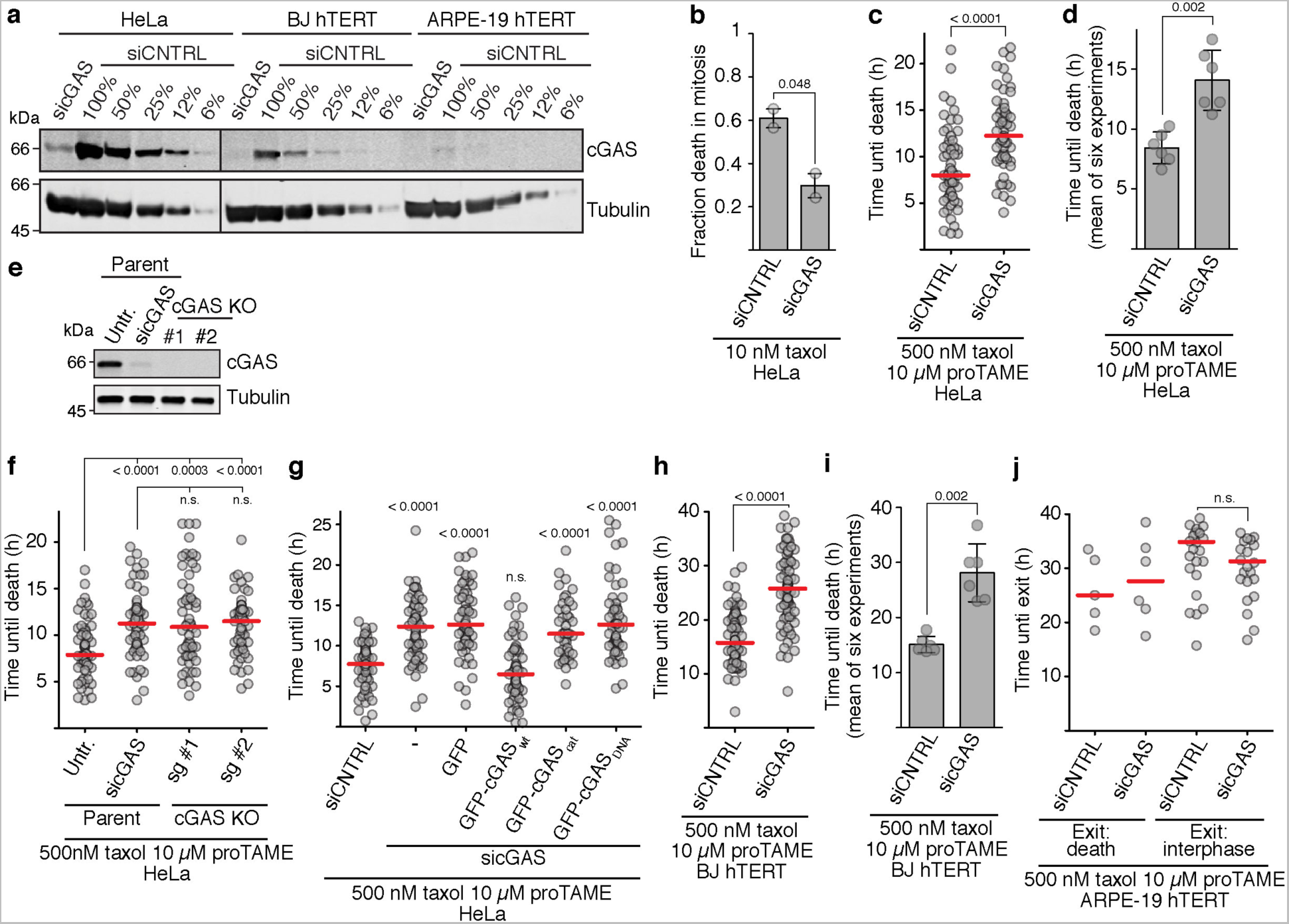
cGAS promotes cell death in mitosis. **a,** Quantitative Western blot analysis of cGAS Dilution series from extracts made from control siRNA treated cells were titrated to assess cGAS depletion. **b-j,** Live microscopy-based analysis of mitotic cell death. Cells were released from G2 arrest into M phase with indicated drugs, and duration of mitosis until mitotic cell death was monitored. All red lines indicate the median. **b,** Fraction of mitotic cell death in HeLa cells, treated with control siRNA (siCNTRL), or cGAS-targeting siRNA (sicGAS). Mean values and range (error bars) from two experiments (n = 100 each) are shown. **c,** Timing of mitotic cell death of individual HeLa cells (grey dots, n = 60 each). **d,** Fraction of mitotic cell death in HeLa cells, treated with control siRNA (siCNTRL), or cGAS-targeting siRNA (sicGAS). Mean values and range (error bars) from two experiments (n = 100 each) are shown. **e, f,** Analysis of HeLa cells treated with siRNA, and of two clones derived using different short guide RNAs. **e**, Western blot analysis of cGAS levels. **f,** Time until mitotic cell death. Grey dots represent individual cells (n = 60 each). sg, small guide RNA. **g,** Complementation analysis in HeLa cells. Timing of mitotic cell death of individual HeLa cells (grey dots, n = ~50 each) was analysed. cGAScat, catalytic mutant. cGASDNA, DNA binding-induced catalytic activity mutant. See Extended Data Fig. 6c for Western blot analysis**. h,** Timing of mitotic cell death of individual BJ-hTERT cells (grey dots, n = 80 each). **i,** Means and standard deviation of timing of death from six experiments of BJ-hTERT cells (grey dots are medians from each individual experiment). **j,** Timing of mitotic cell death of individual ARPE-19 hTERT cells (grey dots, n = 40 each).

Depletion of cGAS in HeLa cells (Fig. 3c, d; Supplementary Video 1; also see Fig. 3e-g) or BJ-hTERT cells (Fig. 3a, h, i) substantially extended mitotic lifespan when compared to non-targeting control siRNA-treated cells. Similarly, HeLa clones harbouring CRISPR-Cas9 mediated disruptions of cGAS generated with two different short guide RNAs showed phenotypes identical to cGAS knockdown (Fig. 3e, f). The extension of mitotic lifespan by cGAS depletion in HeLa cells was comparable to the effect observed upon inhibition of apoptosis with the caspase inhibitor Z-VAD-FMK (Extended Data Fig. 6a). cGAS depletion also greatly extended mitotic lifespan in another HeLa cell line (CCL-2), which had an overall longer mitotic lifespan than the HeLa Flp-In T-REx line (Extended Data Fig. 6b). In contrast, ARPE-19 hTERT retinal pigment epithelium cells, which express very little cGAS (Fig. 3a), tolerated mitotic arrest extremely well, surviving extensive arrest (30 – 40 h), and cGAS siRNA treatment did not have any impact on mitotic lifespan (Fig. 3j).

The extended lifespan in taxol was not due to off-target effects, as expression of siRNA resistant wild type cGAS (GFP-cGAS_wt_) fully restored shorter lifespans in sicGAS cells (Fig. 3g; Extended Data Fig. 6c). Treatment with control siRNA by itself also did not affect timing of death (Extended Data Fig. 6d). Promotion of mitotic cell death requires catalytic activity of cGAS since neither of two catalytic mutants, GFP-cGAS_cat_ (E225A D227A), mutated in the catalytic pocket, or GFP-cGAS_DNA_ (K407E K411A), in which DNA binding does not induce the conformational changes required for catalysis^15^, rescued cGAS depletion (Fig. 3g; Extended Data Fig. 6c). cGAS does not appear to be a general death accelerator, as its depletion did not affect timing of death after UV irradiation (Extended Data Fig. 7a, b). Altogether, these results indicate that cGAS promotes death during mitotic arrest.

If cGAS promotes death through cGAMP, cGAMP addition should substitute for cGAS and reduce mitotic lifespan. In both HeLa and BJ-hTERT cells depleted for cGAS, this was indeed the case (Fig. 4a, b). However, cGAMP treated cells did not die as fast as controls, most likely due to incomplete intracellular cGAMP delivery. In previous studies investigating cGAMP induction of interferon transcription, membrane transversal was promoted by transfection agents^27^ or chemical membrane permeabilisation^5,30^. These treatments are cytotoxic, and are therefore not appropriate for our assay. Regardless, our experiments show that cGAS, at least in part through cGAMP, induces mitotic death in both untransformed and transformed cells. cGAMP-treatment does not appear to induce cell death by a nonspecific mechanism, since cGAMP addition or ectopic cGAS expression in ARPE-19 hTERT cells, which hardly express cGAS (Fig. 3a), had only modest effects on mitotic death (Extended Data Fig. 8a-c). Despite the presence of the cGAMP acceptor STING (Extended Data Fig. 5a), ARPE-19 hTERT cells appear to be defective further downstream in the pathway, as cGAMP addition could not induce IRF3ph (Extended Data Fig. 8d). Altogether, these observations suggest that cGAMP signalling, but not unspecific cGAMP toxicity, promotes mitotic cell death.

**Figure 4.**
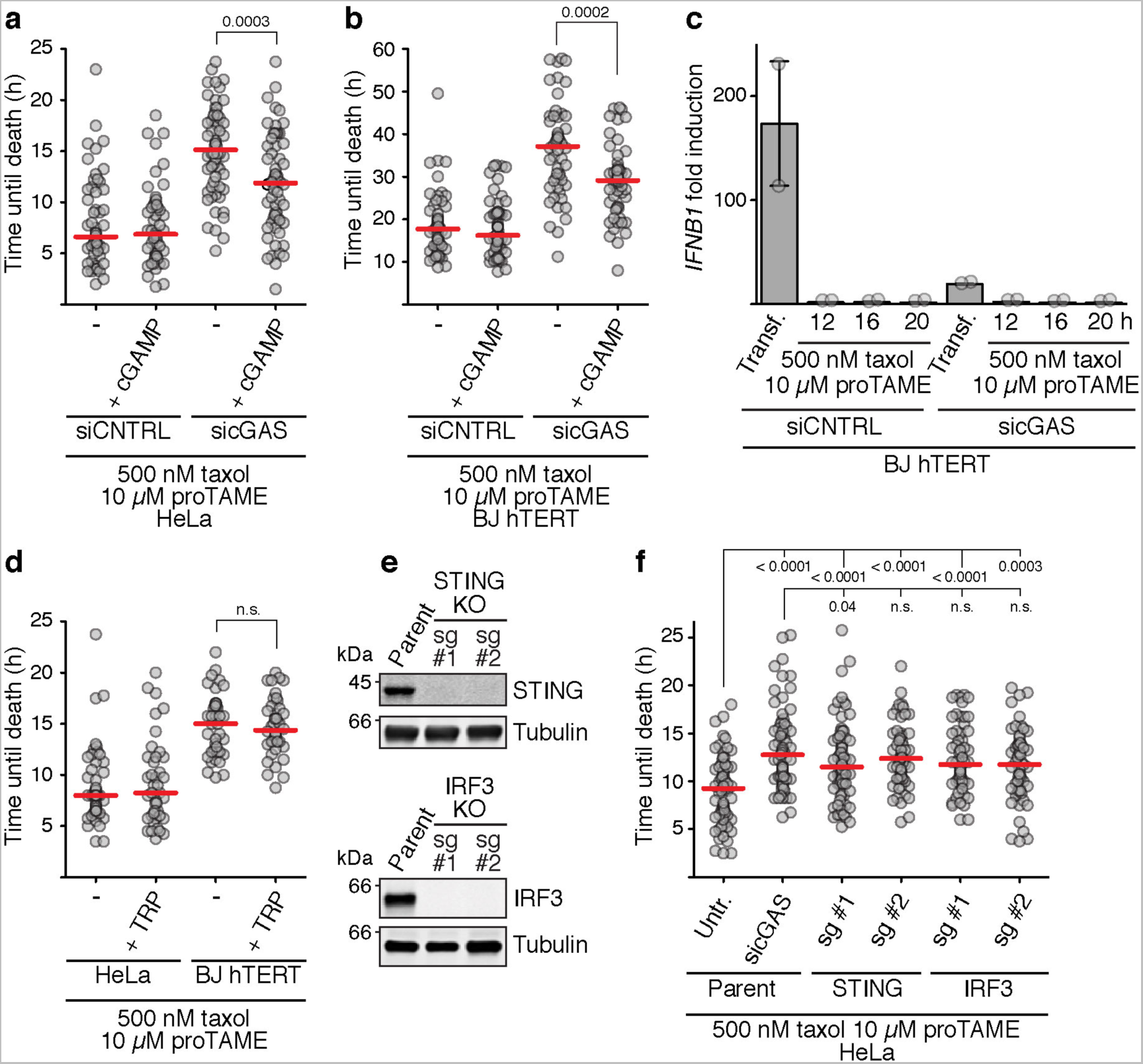
The cGAS-cGAMP-STING-IRF3 axis promotes mitotic cell death independently of transcriptional induction. All red lines indicate the median. siCNTRL, control siRNA. **a and b,** Cells were arrested in G2 and released in the presence of the indicated drugs. After 1.5 h (a) or 4 hours (b), cGAMP was added to 33 µM for the indicated cells. Individual cells (grey dots, n = 50 each) were tracked using live microscopy. **c,** cDNA prepared from the indicated cells was used to determine IFNB1 expression. Transf., transfected with naked DNA. Mean and range from two experiments are shown. **d,** Cells were arrested in G2 and released in the presence of the indicated drugs. After 1.5 h (HeLa) or 4 h (BJ hTERT), triptolide (TRP) was added to 10 µM for the indicated cells. Individual cells (grey dots, n = 60 each) were tracked using live microscopy. **e,** Western blot analysis of STING and IRF3 knockout clones generated with two different short guide RNAs (sg) each. **f,** Live microscopy analysis of cells from the clones shown in (e) released from G2 arrest in the presence of the indicated drugs. Individual cells (grey dots, n = 50 each) were tracked using live microscopy. sg, small guide RNA.

Since the major known consequence of cGAS activation is stimulation of transcription factors, such as IRF3, we wondered how cGAS stimulates cell death in mitosis, where transcription is generally shut off^31^. Indeed, interferon beta, the classical IRF3 target^1^ was not induced during mitosis in BJ-hTERT cells, whilst DNA transfection in asynchronous cells effectively stimulated its transcription (Fig. 4c). Furthermore, treatment of cells with the transcription inhibitor triptolide^32^ did not extend mitotic lifespan (Fig. 4d). However, IRF3 can also induce apoptosis independently of transcription by mediating the mitochondrial translocation of Bax^2^, a protein that induces outer mitochondrial membrane permeabilisation, a critical step for apoptosis in mitosis^22,23,33^. In agreement with the requirement for signalling to IRF3 in the transcription-independent induction of apoptosis, CRISPR-Cas9 mediated disruption of either STING or IRF3 extended mitotic lifespan similarly to cGAS depletion (Fig. 4e, f). Thus, our data show that cGAS induces mitotic cell death through cGAS signalling, but independently of transcription.

Here we demonstrated that cGAS signalling is limited by nucleosomes, suppressing cGAS pathway activation during normal mitosis. However, during defective mitosis, cGAS can be utilised to sense the prolonged exposure of chromosomes to the cytoplasm, promoting cell death. Alternatively, cGAS may become activated by byproducts of extended mitotic arrest, such as mitochondrial permeabilisation^34^ or DNA damage^35,36^, although these events were seen at much later timing (>12 h) than the initial emergence of IRF3 phosphorylation (<1h, Fig. 2h). Ultimately, promotion of cell death by cGAS is likely to occur by stimulating mitochondrial relocalisation of IRF3, which in turn activates Bax-mediated apoptosis^2^. Rather than solely determining the timing of mitotic death, cGAS is likely to collaborate with other independent death factors operating on their own timers. These may include the mitosis-specific degradation of the pro-survival protein Mcl1, as well as transcriptional regulation of pro-apoptotic and pro-survival genes^22,23^. As taxol prolongs mitosis, and also induces formation of micronuclei, where cGAS strongly accumulates, it is tempting to speculate that taxol efficacy may be altered by the functionality of the cGAS pathway. Induction of the intrinsic cell death pathway by mitotic defects may prevent the formation of aneuploidy, frequently associated with cellular senescence and cancer^37–39^. Activation of cGAS by multiple products of defective mitosis such as micronuclei, could also induce inflammation, which can affect tumour development and metastasis, as well as the immunological response to tumour development^40^. A detailed knowledge of cGAS function in the response to mitotic defects will be important for our understanding of tumour development and chemotherapy response.

## Methods

**Antibodies and reagents.** cGAS was detected with D1D3G (# 15102) from Cell Signaling Technology (1:500 dilution for western blot, 1:400 dilution for immunofluorescence); STING was detected with D2P2F (# 13647) from Cell Signaling Technology (1:500 dilution for western blotting); IRF3 (S396ph) was detected with # 4947 from Cell Signaling Technology (1:500 dilution for western blotting); IRF3 (total) was detected with ab68481 (abcam; 1:500 dilution for western blotting); α tubulin was detected with T9026 from Sigma (1:15000 dilution for western blotting). For western blotting, IRDye 800CW and 680LT secondary antibodies were used at 50 ng/ml and detected on an Odyssey infrared imaging system (LI-COR). For immunofluorescence microscopy, Alexa-488 and Alexa-555 coupled secondary antibodies were used (Jackson ImmunoResearch).

cGAMP was obtained from Invivogen (tlrl-nacga23); doxycycline was obtained from Sigma (D9891); proTAME was obtained from Boston Biochem (I-440); RO3306 was obtained from Sigma (#SML-0569); SiR-DNA was obtained from Cytoskeleton (CY-SC007); taxol was obtained from Calbiochem (# 580555); thymidine was obtained from Sigma (T1895). Triptolide was obtained from Invivogen (# ant-tpl).

cGAS siRNA (# L-015607-02, ON-TARGET plus SMARTpool) was obtained from Dharmacon. Non-targeting control siRNA (# D-001810-01, ON-TARGET plus) was obtained from Dharmacon.

**Cell lines, cell culture, siRNA knockdown and cell cycle synchronizations.** The identity of all cell lines was validated using polymorphic short tandem repeat loci (Memorial Sloan Kettering Cancer Center). Mycoplasma contamination was monitored by fluorescence microscopy. HeLa Flp-In T-Rex cells stably expressing H2B-RFP (referred to as HeLa cells throughout this manuscript, a gift of A. Desai and R. Gassmann)^18^, HeLA CCL-2 cells (a gift from I. Nakagawa), BT474, BT549, CAL51, MDA-MB-157, MDA-MB-231 and T47D cells (gifts of K. Birsoy) were grown in DMEM (Gibco 11995065) supplemented with 10% tet-tested FBS (Atlanta Biologicals) and penicillin-streptomycin (100 u/ml, Gibco). HCC1143 cells (a gift of K. Birsoy) were grown in RPMI 1640 medium (Life Technologies SH30027FS) supplemented with 10% tet-tested FBS (Atlanta Biologicals) and penicillin-streptomycin (100 u/ml, Gibco). BJ hTERT cells (a gift of T. de Lange) were grown in DMEM (Gibco 11995065) supplemented with 10% tet-tested FBS (Atlanta Biologicals), penicillin-streptomycin (100 u/ml, Gibco) and 10% medium 199 (Sigma M4530). ARPE-19 hTERT cells (a gift of T. de Lange) were grown in DMEM-F12 (Life Technologies 10565-018) supplemented with 10% tet-tested FBS (Atlanta Biologicals) and penicillin-streptomycin (100 u/ml, Gibco).

HeLa Cell lines expressing GFP-tagged versions of cGAS or IRF3 were generated using plasmid pcDNA5/FRT/TO as a backbone following instructions based on the Flp-In system (Invitrogen). ARPE-19 hTERT cell lines expressing GFP-tagged versions of cGAS were generated using the piggyBac transposon system. cGAS_cat_ represents the E225A D225A double mutant, cGAS_DNA_ represents the K407E K411A double mutant, and the cGAS KKKR mutant is the K407E K411A K173E R176E quadruple mutant. Plasmid construction details are available on request.

siRNA knockdowns were performed using the reverse transfection procedure. Briefly, siRNA was mixed with media without FBS (Opti-MEM, Life Technologies # 51985-034) and Lipofectamine RNAiMAX (Invitrogen # 13778075) at an siRNA concentration of 120 nM (10 nM final concentration once cells and media were added). Lipofectamine RNAiMAX was added at 1.2 µl/ml of final volume. The mixture was incubated for 15 min, and cells were then added in media without antibiotics. Media was changed the next day.

For cell synchronizations, cells were pre-synchronized in S phase for 18 h with 2.5 mM thymidine. Cells were then washed twice in PBS and once in media, and cells were then released from the S phase arrest for 2 h in media without drugs. RO3306 (9 µM final) was then added to capture cells in G2. After 8-10 h, cells were washed twice with PBS, once with media containing 500 nM taxol before further incubation in media containing 500 nM taxol and 10 µM proTAME. If BJ hTERT or ARPE-19 hTERT cells were prepared for time-lapse microscopy, SiR-DNA (125 nM final concentration) was added to the media 2 h before release from G2 arrest, and to the final media containing taxol and proTAME to allow for visualization of DNA. For induction of GFP-IRF3 or GFP-cGAS versions, doxycycline was added to a final concentration of 100 ng/ml ~24 h before release from G2 arrest.

If cells were prepared for time-lapse imaging, cells were subjected to siRNA knockdown at the same time as they were plated at ~30% confluence onto glass bottom culture dishes (MatTek P35GC-1.5-14-C). Cells were grown over night before addition of thymidine. To limit phototoxicity, the final imaging media did not contain phenol red.

**DNA transfections to induce cGAS signalling**. Cells were grown close to confluence on 6 cm dishes. Media was removed from cells and replaced with 2.4 ml of media containing no antibiotics. To generate the transfection mix, 7.5 µg of pAS696^41^ were combined with 0.3 ml Opti-MEM (Life Technologies # 51985-034). Separately, 25 µl Lipofectamine 2000 (Life Technologies, # 11668) were mixed with 0.3 ml Opti-MEM. Both mixtures were incubated at room temperature for 5 min, combined and incubated at room temperature for 20 min before being added to the cell dish. Cells were harvested after two hours.

**CRISPR-Cas9 mediated gene disruptions**. Sequences encoding short guide RNAs (sgRNAs) were cloned into pX330^42^, and transfected into cells. Cells were grown for five days, and then plated for single colonies. Multiple candidate clones were picked and tested for gene disruption by western blot. sgRNA sequences were as follows: cGAS sg#1: CTGCCCCCAAGGCTTCCGCA; cGAS sg#2: GGCGCCCCTGGCATTCCGTG; IRF3 sg#1: GGTTGCGTTTAGCAGAGGAC; IRF3 sg#2: ACCTCTCCGGACACCAATGG; STING sg#1: GCTGGGACTGCTGTTAAACG; STING sg#2: GCAAGCATCCAAGTGAAGGG.

**Western blots.** Cells were removed from dishes by trypsin treatment (Life Technologies # 25300-054), washed in PBS and resuspended in 2x sample buffer (125 mM Tris-Cl [pH 6.8 at 22 ºC]; 2% SDS; 0.7 M 2-mercaptoethanol) and boiled for 5 min. Samples were then subjected to water bath sonication, spun down, and sample buffer containing glycerol and bromophenol blue were added to result in a final concentration of 6.25% glycerol and 25 µg/ml bromophenol blue. Samples were again boiled and subjected to gel electrophoresis and western blotting.

For checking siRNA efficiency, cells treated identically to the cells that were used for microscopy, were harvested just before release into taxol.

**Immunofluorescence microscopy.** Cells were grown on coverslips coated in poly-D-lysine (Sigma P1024, coverslips were coated by incubation for ~30 min in a 1 mg/ml solution). Coverslips were washed once in PBS, and subsequently cells were fixed with paraformaldehyde (Electron Microscopy Services # RT 157-8; 2% in PBS) by incubation for 20 min. Coverslips were washed three times in PBS and cells were then permeabilised for 5 min by incubation with PBS containing 0.2% Triton X-100 (Bio-Rad). Following three washes in PBS, coverslips were blocked in PBS-T (PBS plus 0.1% Tween-20 (Bio-Rad)) containing 5% BSA (Sigma A7906).

Antibody incubations (1 h each) were performed in the same solution. Coverslips were washed five times in PBS-T after each antibody incubation and subsequently mounted in Vectashield with DAPI (Vector Laboratories; H-1200). Microscopy was performed using a Delta Vision Spectris setup (Applied Precision) consisting of an Olympus IX71 wide-field inverted fluorescence microscope, and a Photometrics CoolSnap HQ camera (Roper Scientific). Images were captured at 0.2 µm Z steps, processed by iterative constrained deconvolution using SoftWoRx (Applied Precision), and analysed in ImageJ (v. 2.0.0-rc-43/1.51k). Sum projections are shown throughout the figures.

**Time-lapse microscopy.** Live cell imaging was performed in an LCV110U VivaView FL incubator microscope (Olympus) at 37 °C in 5% CO_2_. The setup also consisted of an X-Cite-exacte illumination source (Excelitas Technologies) and an Orca-R2 CCD camera (Hamatsu). Images were acquired with a 20 × objective every 15 min for 48-72 h, in some cases with multiple Z steps (~2 µM step sizes; ImageJ was used to generate sum projections when Z slices were taken). Individual cells were manually tracked using the CellCognition software^43^ (v 1.5.2). Timing of death or slippage into interphase were determined morphologically (slippage was defined by cell flattening, and initiation of death was defined by the characteristic morphological changes reminiscing of cell explosion). When cells expressing GFP-cGAS or its mutated versions, were used, only cells with similar levels of GFP-cGAS were analysed in order to compensate for expression level heterogeneity within the population.

**UV-viability assays.** Cells were seeded onto 6-well plates at ~20% confluence and allowed to attach for ~10 h. Cells were then washed twice in PBS and irradiated using an XL-1000 Spectrolinker UV crosslinker (Spectronics Corporation). Media was added and cells were grown under standard conditions. At the indicated time points, the supernatant (enriched for dead cells) was transferred to a tube and washed once in PBS, with the wash also moved to the same tube. The remaining cells were trypsinised and also moved to the same tube. Cells were spun down, resuspended in 200 µl PBS and mixed with 50 µl 0.4% Trypan blue. The fraction of dead cells was then determined using cell counting.

**cDNA preparation and quantitative PCR**. cDNA was prepared as follows. ~ 3 × 10^6^ cells were harvested and resuspended in 1 ml TRIzol (Life Technologies, # 15596018). This mixture was frozen at −80 ºC until use. For RNA extraction, 0.2 ml chloroform were added after thawing, and samples were thoroughly mixed. Samples were allowed to stand at room temperature for 2 min before centrifugation for 18 min (10.000 g; 4 ºC). The aqueous phase was removed and mixed with an equal volume of ethanol. After mixing, samples were loaded onto RNeasy columns (Qiagen), washed once with buffer RW1 and twice with buffer RPE. Samples were eluted using 50 µl of nuclease-free water. ~ 300 ng RNA of each sample were used to generate cDNA using the Transcriptor First Strand cDNA Synthesis kit (Roche, # 04 379 012 001) according to the manufacturer’s instructions (both Anchored-oligo (dT)_18_ as well as random hexamer primers were used).

2 µl of each cDNA reaction were used for quantitative PCR with the LiqhtCycler 480 SYBR Green I Master mix (Roche, # 04 707 516 001), carried out on an iCycler (Bio-Rad) equipped with the IQ 5 detection system. Interferon beta (*IFNB1*) primer sequences were AGGACAGGATGAACTTTGAC and TGATAGACATTAGCCAGGAG. Readings were normalized to *GAPDH* using primers AATCCCATCACCATCTTCCA and TGGACTCCACGACGTACTCA. The following temperature program was used: 5 min 95 ºC; 50 × (10 sec 95 ºC; 20 sec 52 ºC; 20 sec 72 ºC). For plotting, signal was normalised to cells synchronised in G2 but otherwise untreated.

**Recombinant proteins.** Nucleosomes were prepared from pre-formed H2A–H2B dimers and H3–H4 tetramers. Our previously published methods were used for histone purification^13^. Briefly, histones H2A, H2B, H3 and H4 were individually expressed in *E. coli* under conditions in which they formed inclusion bodies. Histones were then individually purified under denaturing conditions (6 M Guanidine HCl; 500 mM NaCl; 50 mM Tris-Cl [pH 8 at 22 ºC]; 5 mM 2-mercaptoethanol; 7.5 mM imidazole), refolded into H2A–H2B dimers or H3–H4 tetramers by dialysis into 20 mM MOPS; 500 mM NaCl; 2% glycerol; 1 mM EDTA; 5 mM 2-mercaptoethanol; pH 7 at 22ºC, and purified using size-exclusion chromatography.

cGAS was purified using a modified version of published procedures^9^. Similar to previous work^8,9,44^, we generated cGAS lacking its N terminal (amino acids 1-150) unstructured tail. This truncation had previously been shown to be catalytically active^8,9,44^, and to fully substitute for full-length cGAS in interferon induction in response to cytoplasmic DNA^4^. We expressed this protein in E. coli with an N-terminal MBP tag and a C-terminal 10xHis tag, with the tags separated from the rest of the protein with TEV protease recognition sites. *E. coli* Rosetta 2 (DE3 pLysS) cells containing cGAS expression plasmids were grown in 1.5 × TBG-M9 medium (15 g/l tryptone; 7.5 g/l yeast extract; 5 g/l NaCl; 0.15 g/l MgSO_4_; 1.5 g/l NH_4_Cl; 3 g/l KH_2_PO_4_; 6 g/l Na_2_HPO_4_; 0.4 % glucose) until an OD_600_ of ~0.9, and protein expression was induced at 18 °C with 0.6 mM IPTG for 20 h. Cells were then lysed by sonication in wash/lysis buffer (50 mM Tris-Cl [pH 7.5 at 4 °C]; 0.3 M NaCl; 10 mM 2-mercaptoethanol) supplemented with 1 mM PMSF; 0.25 mg/ml lysozyme; 10 µg/ml leupeptin; 10 µg/ml pepstatin; 10 µg/ml chymostatin. The fusion protein was enriched on amylose resin (NEB), and eluted by treatment with TEV protease. To remove TEV protease (which itself was His-tagged), the eluate was also run over a Ni-NTA column (Qiagen). Proteins were further purified on a HiTrap heparin column (GE Healthcare) on an ÄKTA FPLC system (GE Healthcare) using a salt gradient for elution that was established with wash/lysis buffer as low salt buffer and wash/lysis buffer containing 1 M NaCl as high salt buffer. Peak fractions were subjected to a final purification step by size-exclusion chromatography using a Superdex 200 10/300 GL column (GE Healthcare) on an ÄKTA FPLC system (GE Healthcare) in 20 mM Tris-Cl [pH 7.5 at 4 °C]; 0.3 M NaCl; 1 mM DTT. Peak fractions were combined, concentrated, and stored at −20 °C in the same buffer containing 48% glycerol.

**Preparation of nucleosome arrays and mononucleosomes for binding assays.** DNA substrates used for nucleosome arrays and for their naked DNA counterparts were generated as described previously^13,41^. Briefly, plasmid pAS696 containing 19 tandem copies of the “601” nucleosome-positioning sequence^45^ was digested with HaeII, DraI, EcoRI and XbaI to remove the 601 array and to digest the vector backbone into smaller pieces. The array was then separated from the vector backbone fragments using PEG precipitation. To facilitate coupling to streptavidin beads, ends were filled in with Klenow fragment (NEB) in the presence of dCTP, biotin-14-dATP (Invitrogen), thio-dTTP and thio-dGTP (both Chemcyte). This resulted in biotinylation of both ends of the linear array. Arrays were separated from unincorporated nucleotides using Sephadex G-50 Nick columns (GE Healthcare).

To generate the DNA used for mononucleosomes or single copies of the naked 601 sequence, pAS696 was digested with AvaI (which cuts in between 601 monomers), and the fragment was separated from the vector backbone using PEG precipitation.

Nucleosomes were assembled on these substrates using salt dialysis following published procedures^13,41^. 50 µl reactions were prepared containing 10 µg of DNA, and histones H2A– H2B and H3–H4 at empirically determined concentrations that varied slightly with each batch (but usually in slight excess over DNA), 2 M NaCl and 1x TE. Mixtures were moved to dialysis buttons (Hampton Research), submerged in 500 ml high salt buffer (10 mM Tris-Cl [pH 7 at 22 °C]; 1 mM EDTA; 2 M NaCl; 5 mM 2-mercaptoethanol; 0.01 % Triton X-100) and salt was reduced through dialysis over an exponential gradient established by pumping in 2 l of low salt buffer 1 (as high salt buffer but 50 mM NaCl) whilst removing excess buffer from the original 500 ml over 1-3 days at 4 °C. Nucleosomes were then dialyzed into low salt buffer 2 (10 mM Tris-Cl [pH 7 at 22 °C]; 0.25 mM EDTA; 100 mM NaCl; 1 mM TCEP). Mononucleosomes that were used for cGAMP synthesis assays were further dialyzed into 20 mM Tris (pH 7.5 at 22 °C), 150 mM NaCl, 1 mM DTT, 10 mM MgCl_2_. The quality of nucleosome formation on arrays was assessed by digestion with AvaI (which cuts in between 601 monomers on the array) followed by native gel electrophoresis in 5% polyacrylamide using 0.5x TBE as running buffer, as described previously^41^. Quality of mononucleosome formation was determined by native gel electrophoresis without prior digestion.

**Generation of beads coupled to naked DNA or nucleosome arrays.** For both nucleosome beads and naked DNA beads, nucleosomes were initially coupled to Dynabeads M280 Streptavidin (Invitrogen). Nucleosomes containing a total amount of 330 ng of DNA were coupled to 2 µl beads in 75 µl of buffer containing 2.5 % polyvinylalcohol, 150 mM NaCl, 50 mM Tris-Cl (pH8 at 22 ºC), 0.25 mM EDTA, 0.05 % Triton X-100. Coupling was carried out under agitation at room temperature for 2 h. Beads were then washed four times in low salt wash buffer 150 mM NaCl, 50 mM Tris-Cl (pH8 at 22 ºC), 0.25 mM EDTA, 0.05 % Triton X-100. For naked DNA beads, nucleosomes were subsequently stripped by incubation under agitation at 20 ºC in 2 M NaCl, 50 mM Tris-Cl (pH8 at 22 ºC), 0.25 mM EDTA, 0.05 % Triton X-100 for 10 min. Beads were then washed four times in the same buffer, and once in low salt wash buffer.

**Cell-free *Xenopus* egg extracts.** Cytostatic factor (CSF) M phase arrested *X. laevis* egg extracts were prepared as previously described^46^. Histones H3 and H4 were depleted using antibodies recognizing acetylated lysine 12 on histone H4 using our previously describedmethod^13^. The protocol for work with *X. laevis* was approved by The Rockefeller University Institutional Animal Care and Use Committee.

**Bead binding assays.** For bead pull downs in the absence of egg extract, nucleosome beads, chromatin beads and uncoupled beads were washed twice in binding buffer (20 mM Hepes pH 7.7; 200 mM NaCl; 0.05 % Triton X-100; 0.5 mM TCEP pH 7.5).^35^ S labelled full-length cGAS was generated with the TnT Coupled Reticulocyte Lysate System (Promega) according to the manufacturer’s instructions. TnT reactions were diluted 1:10 in binding buffer supplemented with BSA to a final concentration of 0.25 µg/µl, and beads were incubated in 20 µl of this mixture at 20 ºC for 45 min under agitation. Beads were then washed in binding buffer, bound proteins were eluted with SDS sample buffer, and analysed by gel electrophoresis followed by staining with coomassie brilliant blue and fluorography. In parallel, DNA was extracted from another set of beads subjected to an identical procedure to determine whether nucleosome beads and DNA beads contained equal amounts of DNA. To this end, beads were incubated in 500 µl Stop solution 2 (20 mM Tris pH 8.0 at 22 ºC; 20 mM EDTA; 0.5% SDS; 1 mg/ml Proteinase K [Roche]) at 37 ºC for 45 min. DNA was then extracted using phenol/chloroform, precipitated with ethanol/NaOAc in the presence of glycogen, and run on an agarose gel. Bands were visualized using SYBR-safe (Invitrogen), and quantified using ImageJ (v. 2.0.0-rc-43/1.51k).

For bead pull downs from egg extract, nucleosome beads, chromatin beads and uncoupled beads were washed twice with CSF-XB (100 mM KCl; 1 mM MgCl_2_; 50 mM sucrose; 5 mM EGTA; 10 mM Hepes pH 8), and then mixed with 20 µl egg extract containing^35^ S labelled full-length cGAS generated as above (1:10 dilution of TnT reaction in egg extract). Cycloheximide (Sigma) was added to a final concentration of 100 µg/ml to prevent labelling of other proteins within the extract. The mixture was then incubated at 16 ºC under rotation for 45 min, diluted 10 fold in CSF-XB, and beads were recovered. Beads were washed 3 times in CSF-XB containing 0.05% Triton X-100. Bound proteins were eluted with SDS sample buffer and analysed by gel electrophoresis. The gel was stained with coomassie brilliant blue and cGAS was detected with fluorography. Control beads to determine loading amounts were prepared as above with the following exceptions: Mixtures were diluted 10 fold in CSF-XB, beads were recovered, resuspended in 250µl Stop buffer 1 (20 mM Tris pH 8.0 at 22 ºC; 20 mM EDTA; 0.5% SDS; 50 µg/ml RNase A [Qiagen]) and incubated at 37 ºC for 25 min. 250 µl of Stop buffer 2 was then added and beads were incubated at 37 ºC for another 25 min. DNA was extracted with phenol/chloroform, precipitated with ethanol/NaOAc, dissolved in 1x TE containing 50 µg/ml RNase A, and incubated at 37 ºC for 20 min before being analysed as above.

**In-solution binding assays.** Binding reactions contained mononucleosomes or 601 monomers at 20 nM and recombinant cGAS at the concentrations indicated in Fig. 2C in a total of 20 µl of low salt buffer 2 (10 mM Tris-Cl [pH 7 at 22 °C]; 0.25 mM EDTA; 100 mM NaCl; 1 mM TCEP). Reactions were allowed to proceed for 1 h at room temperature. Products were separated by native polyacrylamide gel electrophoresis (5% polyacrylamide in 0.5x TBE). Gels were stained with SYBR-Safe to visualize DNA, and disappearance of the band corresponding to naked DNA or nucleosomes was quantified using ImageJ. Binding affinity was determined with GraphPad Prism (v. 5.0a) using non-linear regression analysis assuming specific one site binding.

**cGAMP synthesis assays.** cGAMP synthesis assays were performed in 20 mM Tris (pH 7.5 at 22 °C), 150 mM NaCl, 1 mM DTT, 10 mM MgCl_2_. Reactions contained 1 µM cGAS and 230 nM mononucleosomes or monomer of naked 601 sequence. cGAS was first pre-incubated with naked DNA or mononucleosomes at room temperature for 30 min. ATP and GTP were then added to a final concentration of 1 mM each to start the reaction. At the same time, GTP labelled at the alpha position with^33^ P was also added to a final concentration of 33 nM. Reactions (10 µl total volume) were incubated at 37 °C and 1.5 µl samples were taken at the indicated time points. Unreacted nucleotides were digested with 4 u alkaline phosphatase (calf intestinal phosphatase, NEB) at 37 °C for 30min, and products were separated via thin-layer chromatography using PEI cellulose F coated sheets (EMD Millipore # 1055790001) with 1.2 M KH_2_PO_4_ (pH 3.8) as running buffer. Plates were dried at 65 °C, analysed by fluorometry, and images were quantified using ImageJ. Initial reaction velocities were determined using linear regression analysis of the 10 min, 20 min and 30 min time points. We confirmed that under our conditions, all cGAS molecules were saturated with nucleosomes or DNA by titration experiments. Thus, the values obtained for our initial reaction velocities represent maximum reaction velocities (Vmax). Vmax values were used to determine Kcat values.

**Statistics.** Statistical significance was determined with GraphPad Prism (v. 5.0a) using a built-in unpaired t-test function (Fig. 2c, d; Fig. 3b; Extended Data Fig. 4b; as determined by F-test, no statistically significant differences in variances existed in any of these cases), or a built-in Mann-Whitney *U* test function (Fig. 3c, d and f-j; Fig. 4a, b, d, f; Extended Data Fig. 6a, b). Since mitotic cell death is a stochastic event, data variations are inherit to each cell line and unpredictable, and we therefore empirically determined the sample size that is sufficient to provide statistically significant differences. No statistical tests were used to predetermine sample sizes. No samples were excluded from any analyses. Investigators were not blinded during experiments and analysis. No randomisation methods were used to determine how samples were allocated.

**Data availability.** The data that support the findings of this study are available from the corresponding author upon reasonable request.

**References**

## Supplementary Information

**Supplementary Video 1.** Time lapse of cGAS-depleted HeLa cells expressing GFP (left) or siRNA resistant GFP-cGAS (right), and H2B-RFP during mitotic arrest in 500 nM taxol and 10 µM proTAME. Top: Overlays of H2B-RFP and GFP (left) or GFP-cGAS (right). Bottom: DIC channel images corresponding to the fluorescence images on top.

## Acknowledgments

We thank K. Birsoy, A. Desai, T. de Lange, R. Gassmann and I. Nakagawa for cell lines, G. Alushin, T. de Lange, C. Jenness, D. Wynne and A. Yoney for critical reading of the manuscript and helpful suggestions; R. Benezra, for discussions and comments on the manuscript, C. Adura Alcaino, K. Birsoy, F. Glickman, L. Lama, U. Schaefer, S., J. Xue and members of the Funabiki lab for helpful discussions; A. Yoney for help with RT-PCR; and members of the Bio-Imaging Resource Center at Rockefeller University for support with imaging. H.F (R01GM075249) and Bio-Imaging Resource Center (S10RR031855) are supported by grants from National Institutes of Health. The content is solely the responsibility of the authors and does not necessarily represent the official views of the NIH.

## Author Contributions

C.Z. and H.F. designed the study. C.Z. carried out all experiments. C.Z. and H.F. wrote the manuscript.

## Author Information

The authors declare that no competing financial interests exist.

## Figures

**Extended Data Figure 1.**
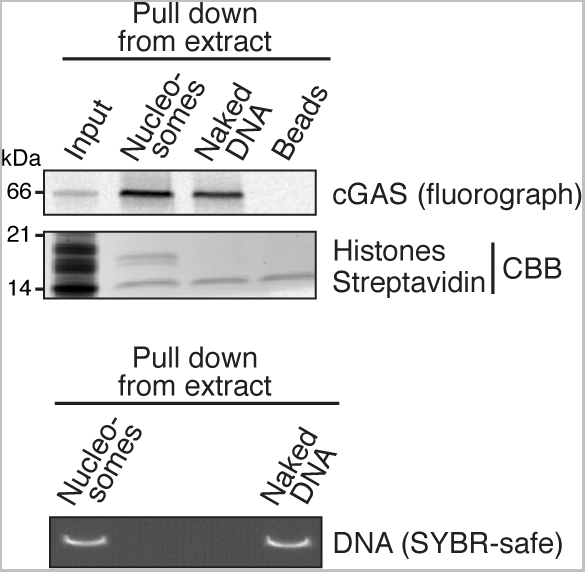
cGAS associates with nucleosomes in M-phase Xenopus egg extracts. The indicated beads were incubated in egg extract containing in vitro generated cGAS. Beads were recovered, washed, and associated proteins were detected by gel electrophoresis followed by fluorometry (upper panel). Note that the extract was depleted of endogenous histones H3–H4 to prevent nucleosome assembly on naked DNA in extract. CBB, Coomassie brilliant blue. Streptavidin serves as an indicator for the amount of beads loaded. The lower panel shows amounts of DNA immobilized on the indicated beads.

**Extended Data Figure 2.**
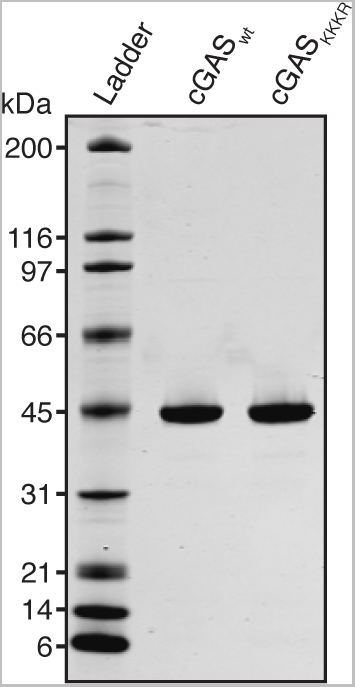
Coomassie stained gel of purified wild type and KKKR mutant cGAS.

**Extended Data Figure 3.**
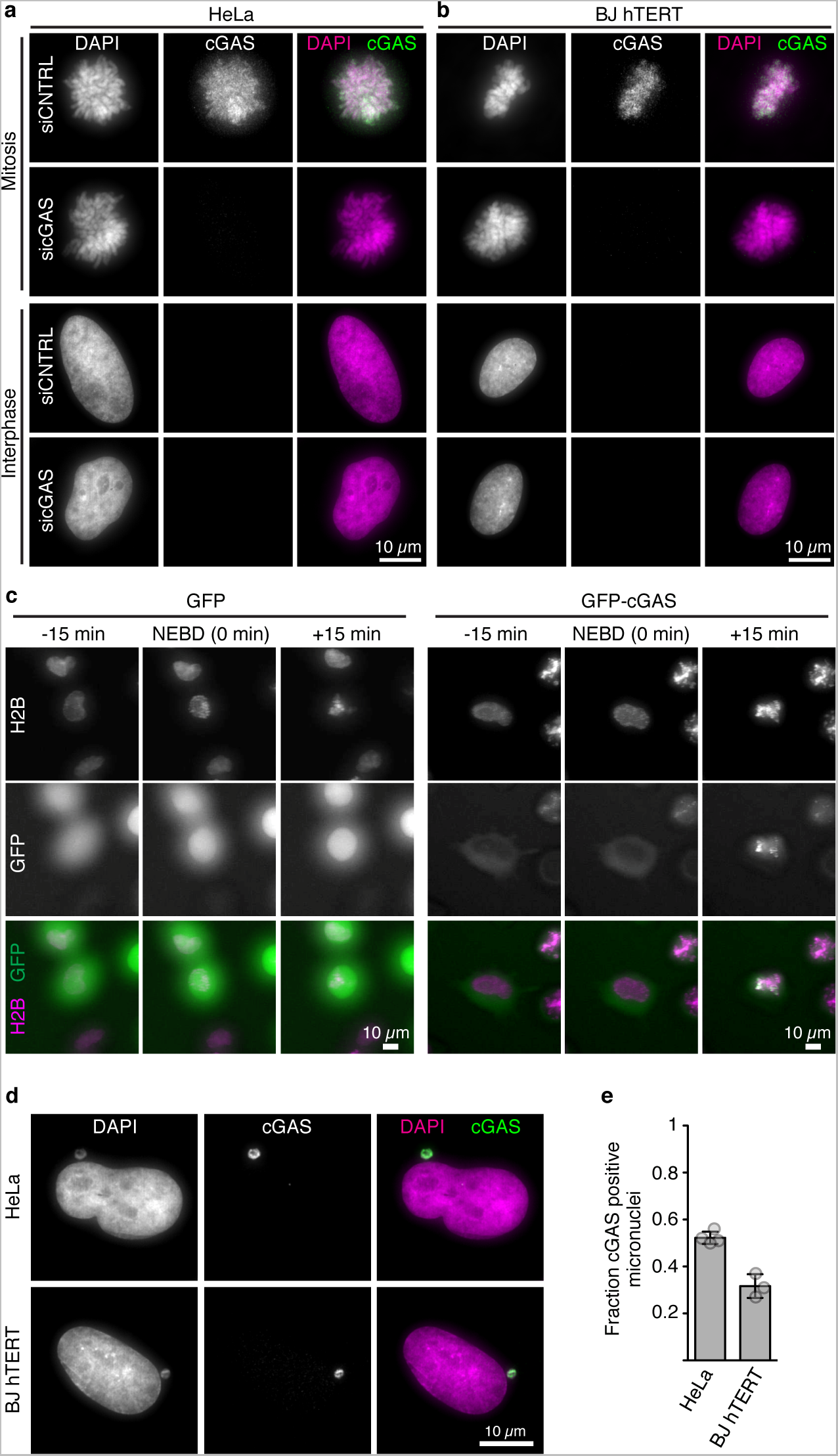
Further analysis of cGAS association with mitotic chromosomes. cGAS signal on mitotic chromosomes is specific. **a** and **b**, cGAS signal on mitotich chromosomes is specific. HeLa cells (a) or BJ hTERT fibroblasts (b) treated with either siRNA to cGAS or with control siRNA (siCNTRL) were stained with anti-cGAS. Note that the siCNTRL cells are the same as the ones shown in Fig. 2e, f. **c**, cGAS associates with mitotic chromosomes in living cells. Live analysis of HeLa cells stably expressing the indicated constructs released from G2 arrest into 500 nM taxol. NEBD, nuclear envelope breakdown. **d** and **e**, cGAS associates witha subset of miconuclei. Immunofluorescence analyis of asynchronous cells from the indicated cell lines stained with anti-cGAS antibodies. **e**, Quantification of (d). Mean values and SEM are shown from three indpendent experiments (n = 100 each).

**Extended Data Figure 4.**
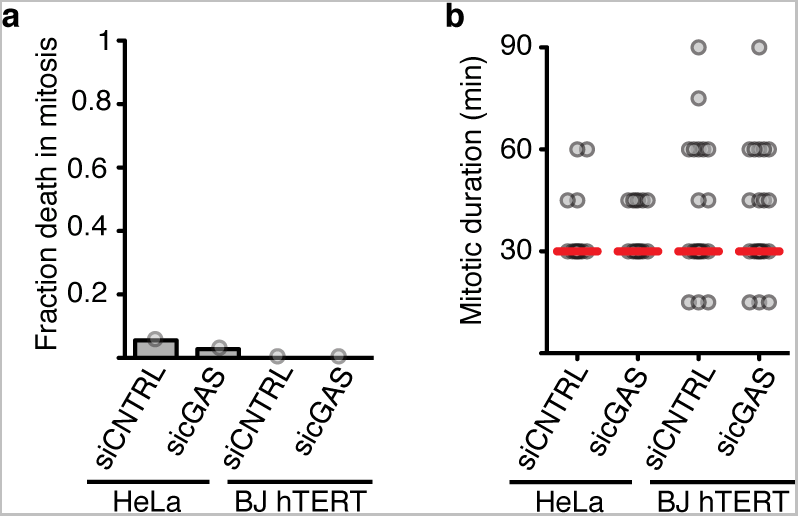
cGAS depletion does not affect unperturbed mitosis. **a**, Fraction of cells of the indicated cell line and treatment that die in an unperturbed mitosis (n = 30 – 50 for each sample). **b**, Median length of mitosis (nuclear envelope breakdown - anaphase) in the indicated cell lines. Red line: median.

**Extended Data Figure 5.**
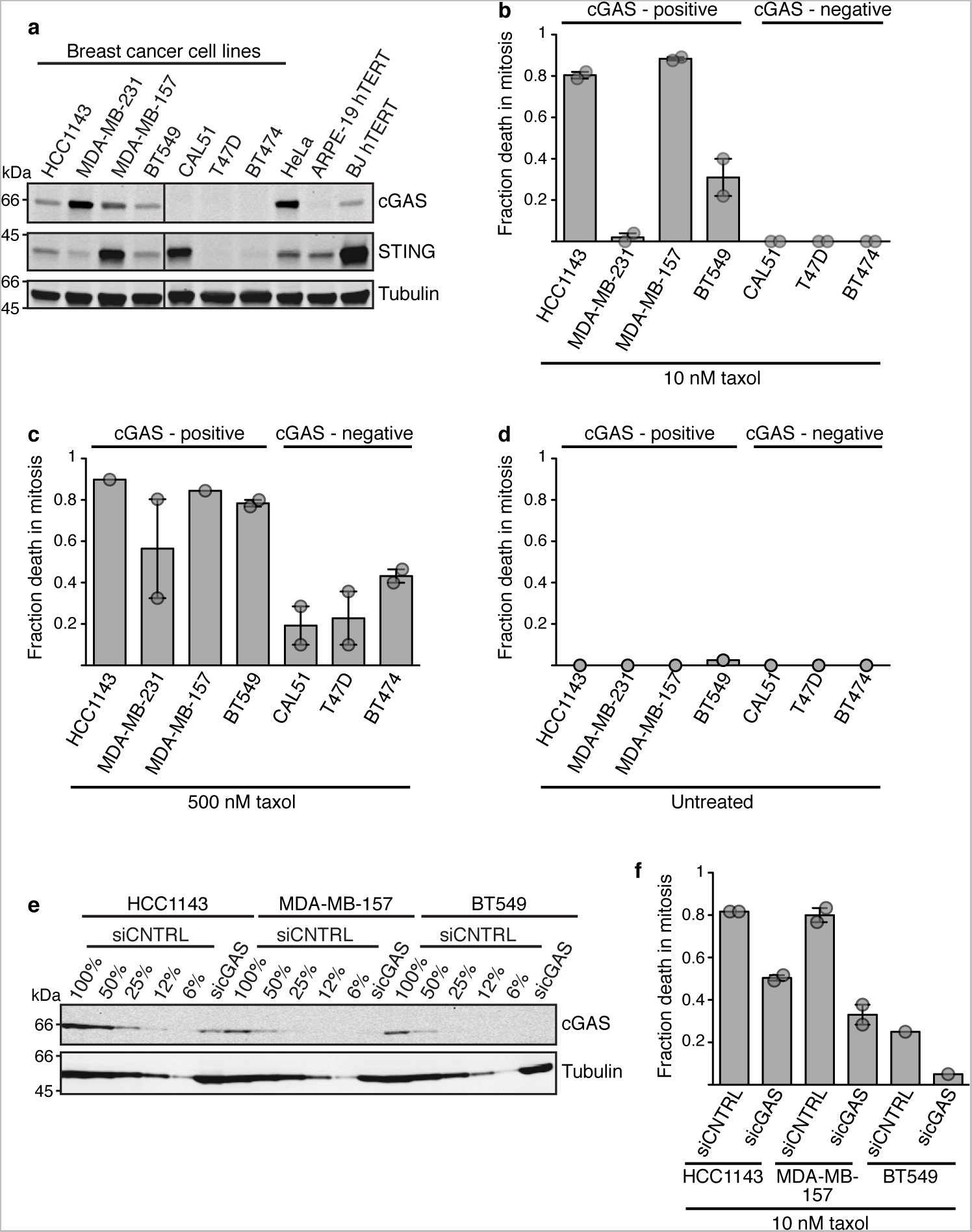
cGAS expression correlates with taxol sensitivity in a panel of breast cancer cell lines. siCNTRL, control siRNA. **a**, Comparison of cGAS and STING expression levels. **b**, Cells were treated with 10 nM taxol, and individual cells were tracked using live microscopy (n = 50 for each sample). Averages and range from two experiments are shown. **c**, Cells were treated with 500 nM taxol, and individual cells were tracked using live microscopy (n = 45 for each sample). Data from one (HCC1143, MDA-MB-157) or two experiments (all others, averages and range are plotted) are shown. **d**, Individual cells grown in the absence of taxol were tracked using live microscopy (n = 40 for each sample). **e**, siRNA-mediated knockdown efficiencies. Percentages indicate dilution series of samples prepared from control siRNA treated cells. **f**, Cells were subjected to cGAS knockdown or control siRNA. Cells were treated with 10 nM taxol, and individual cells were tracked using live microscopy (n = 60 for each sample). Data from one (BT549) or two experiments (HCC1143 and MDA-MB-157, averages and range are plottted) are shown.

**Extended Data Figure 6.**
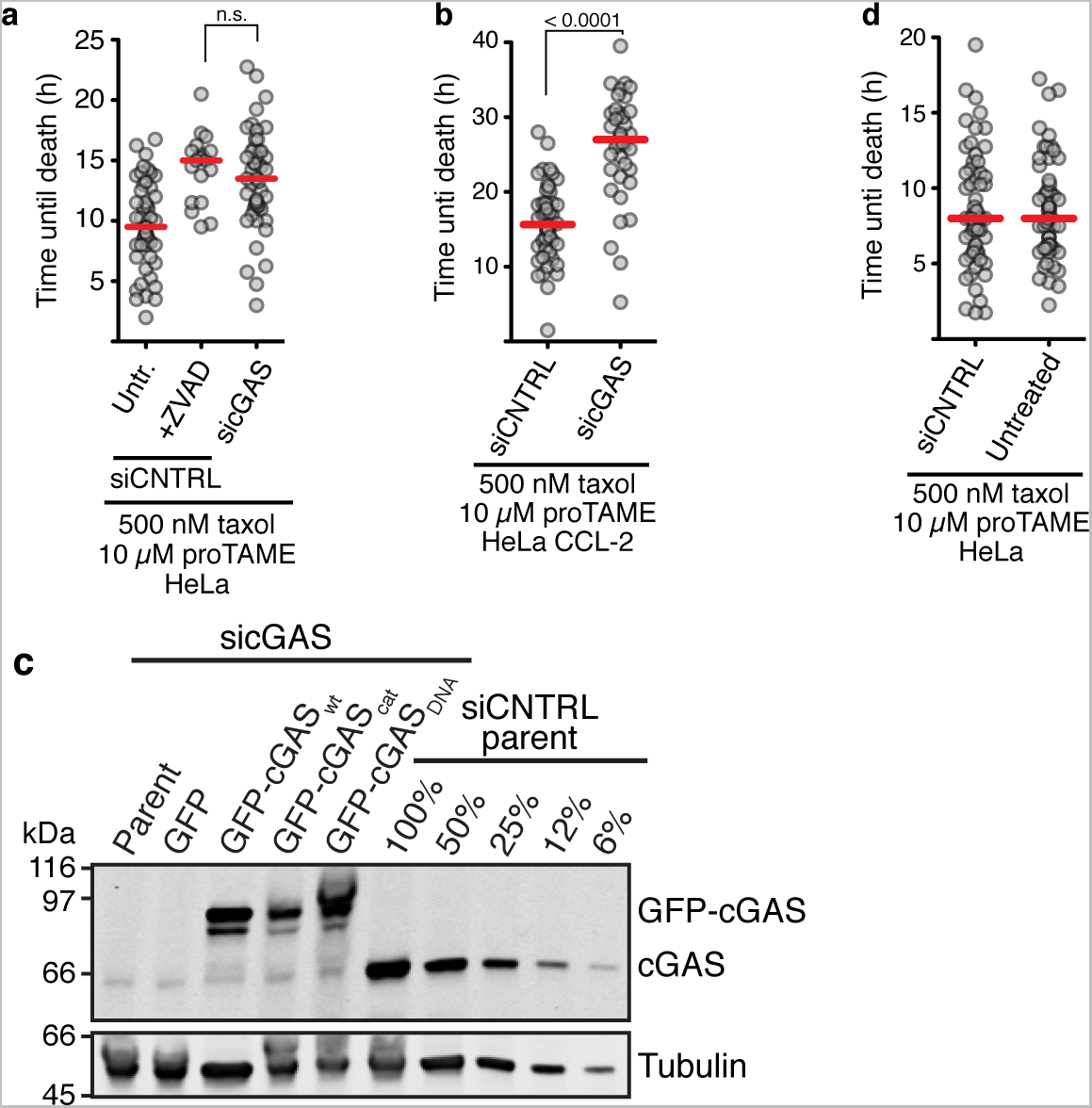
Further characterisation of the effect of cGAS depletion. **a, b, d,** Live microscopy-based analysis of mitotic cell death. Cells were released from G2 arrest into M phase with indicated drugs, and duration of mitosis untill mitotic cell death was monitored. All red lines indicate the median. **a**, The effect of the caspase inhibitor Z-VAD-FMK (ZVAD, 80 µM) was compared with sicGAS. Timing of mitotic cell death of individual HeLa cells (grey dots, n = 50 each) was monitored. **b,** HeLa CCL-2 cells have a longer mitotic lifespan that is further extended by cGAS depletion. Duration of mitosis until mitotic cell death was monitored by live microscopy. Each dot represents an individual cell (n = 40 for each condition). **c**, cGAS expression levels in the experiment shown in Fig. 3g, as determined by western blot analysis. siCNTRL, control siRNA. Percentages indicate dilution series of samples prepared from control siRNA treated cells. cGAS_cat_, catalytic mutant. cGAS_DNA_, DNA binding-induced catalytic activity mutant. **d**, Control siRNA treatment does not affect mitotic cell death. Each dot represents an individual cell (n = 60 for each condition).

**Extended Data Figure 7.**
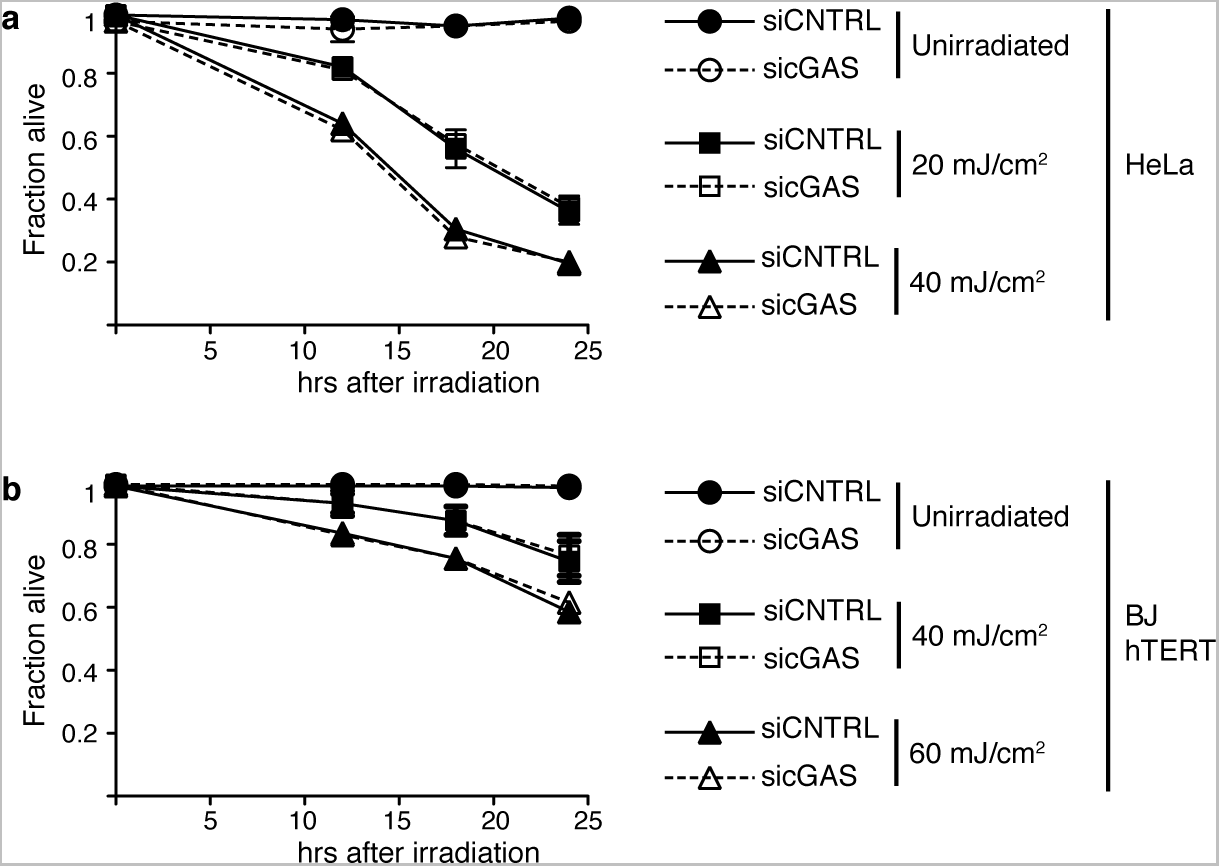
cGAS does not affect survival after UV irradiation. **a** and **b**, Hela cells (**a**) or BJ h-TERT fibroblasts (**b**) were irradiated with the indicated doses of UV, and cell viability was determined at the indicated time points. Mean values and range (error bars) from two experiments are shown. siCNTRL, control siRNA. Note that for many of the data points, error bars are too small to be seen.

**Extended Data Figure 8.**
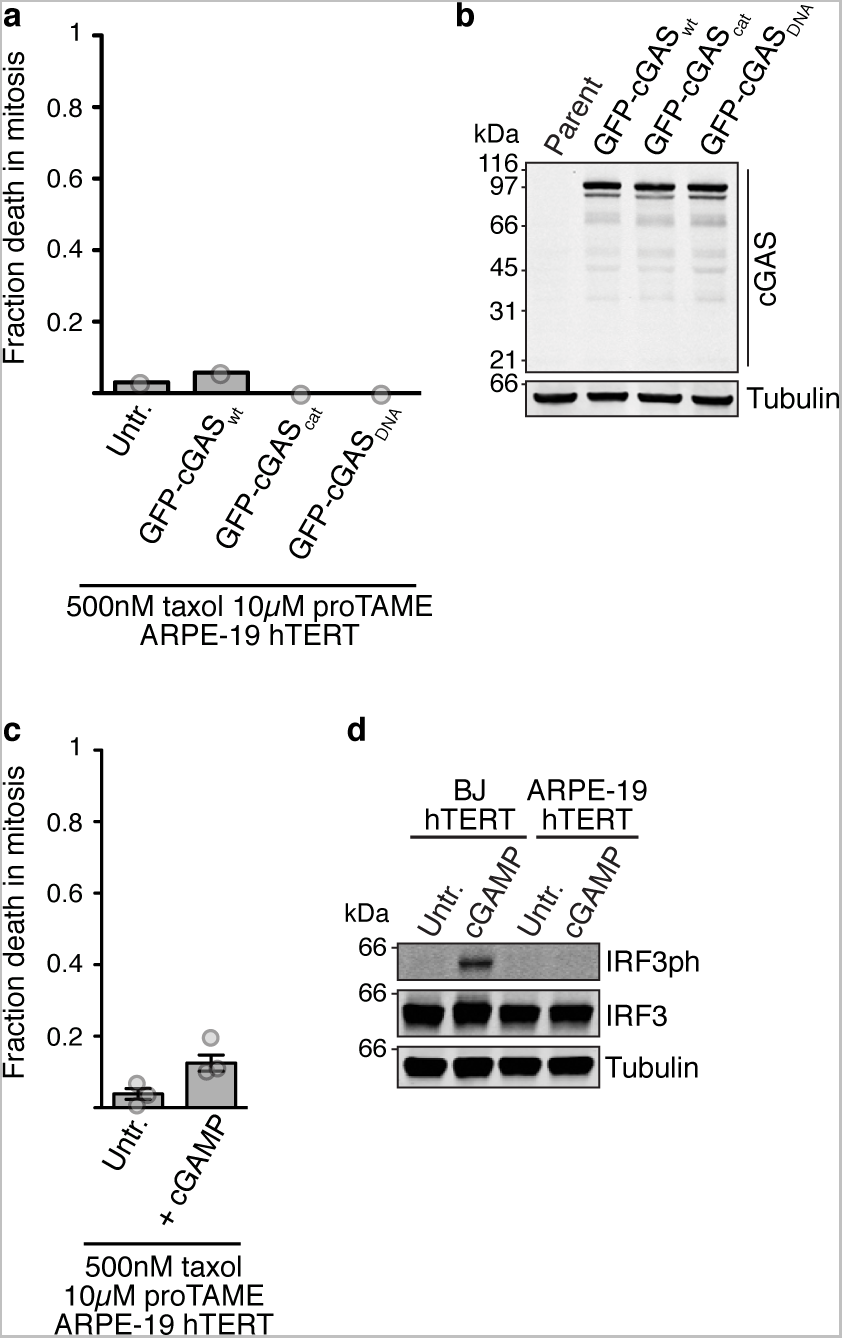
cGAS expression and cGAMP have only minor effect on ARPE-19 hTERT cells. **a** and **b**, ARPE-19 hTERT cells of the indicated type were released from G2 arrest in the presence of taxol and proTAME. Individual cells (n = 30-60 each) were tracked using live-microscopy. **c**, ARPE-19 hTERT cells of the indicated type were released from G2 arrest in the presence of taxol and proTAME. After three hours, cGAMP was added to 33 µM for the indicated cells. Untr., untreated. Mean and SEM from three experiments (n = 30 each) are shown. **d**, Western blot analysis of IRF3 phosphorylation following treatment of asynchronous cells with 33 µM cGAMP.

## References

1. Chen, Q., Sun, L. & Chen, Z. J. Regulation and function of the cGAS-STING pathway of cytosolic DNA sensing. Nat. Immunol. 17, 1142–1149 (2016).

2. Chattopadhyay, S. et al. Viral apoptosis is induced by IRF-3-mediated activation of Bax. EMBO J 29, 1762–1773 (2010).

3. Petrasek, J. et al. STING-IRF3 pathway links endoplasmic reticulum stress with hepatocyte apoptosis in early alcoholic liver disease. Proceedings of the National Academy of Sciences 110, 16544–16549 (2013).

4. Sun, L., Wu, J., Du, F., Chen, X. & Chen, Z. J. Cyclic GMP-AMP synthase is a cytosolic DNA sensor that activates the type I interferon pathway. Science 339, 786–791 (2013).

5. Wu, J. et al. Cyclic GMP-AMP is an endogenous second messenger in innate immune signaling by cytosolic DNA. Science 339, 826–830 (2013).

6. Tang, C.-H. A. et al. Agonist-Mediated Activation of STING Induces Apoptosis in Malignant B Cells. Cancer Res 76, 2137–2152 (2016).

7. Diner, B. A., Lum, K. K., Toettcher, J. E. & Cristea, I. M. Viral DNA Sensors IFI16 and Cyclic GMP-AMP Synthase Possess Distinct Functions in Regulating Viral Gene Expression, Immune Defenses, and Apoptotic Responses during Herpesvirus Infection. MBio 7, e01553–16 (2016).

8. Ablasser, A. et al. cGAS produces a 2‘-5’-linked cyclic dinucleotide second messenger that activates STING. Nature 498, 380–384 (2013).

9. Gao, P. et al. Cyclic [G(2',5')pA(3‘,5’)p] is the metazoan second messenger produced by DNA-activated cyclic GMP-AMP synthase. Cell 153, 1094–1107 (2013).

10. Ishikawa, H. & Barber, G. N. STING is an endoplasmic reticulum adaptor that facilitates innate immune signalling. Nature 455, 674–678 (2008).

11. Raab, M. et al. ESCRT III repairs nuclear envelope ruptures during cell migration to limit DNA damage and cell death. Science 352, 359–362 (2016).

12. Denais, C. M. et al. Nuclear envelope rupture and repair during cancer cell migration. Science 352, 353–358 (2016).

13. Zierhut, C., Jenness, C., Kimura, H. & Funabiki, H. Nucleosomal regulation of chromatin composition and nuclear assembly revealed by histone depletion. Nat Struct Mol Biol 21, 617–625 (2014).

14. Zierhut, C. & Funabiki, H. Nucleosome functions in spindle assembly and nuclear envelope formation. Bioessays 37, 1074–1085 (2015).

15. Civril, F. et al. Structural mechanism of cytosolic DNA sensing by cGAS. Nature 498, 332–337 (2013).

16. Yang, H., Wang, H., Ren, J., Chen, Q. & Chen, Z. J. cGAS is essential for cellular senescence. Proceedings of the National Academy of Sciences 9, 201705499 (2017).

17. Tighe, A., Staples, O. & Taylor, S. Mps1 kinase activity restrains anaphase during an unperturbed mitosis and targets Mad2 to kinetochores. J Cell Biol 181, 893–901 (2008).

18. Gassmann, R. et al. Removal of Spindly from microtubule-attached kinetochores controls spindle checkpoint silencing in human cells. Genes Dev 24, 957–971 (2010).

19. Crasta, K. et al. DNA breaks and chromosome pulverization from errors in mitosis. Nature 482, 53–58 (2012).

20. Hatch, E. M., Fischer, A. H., Deerinck, T. J. & Hetzer, M. W. Catastrophic nuclear envelope collapse in cancer cell micronuclei. Cell 154, 47–60 (2013).

21. Gascoigne, K. E. & Taylor, S. S. Cancer cells display profound intra- and interline variation following prolonged exposure to antimitotic drugs. Cancer Cell 14, 111–122 (2008).

22. Topham, C. H. & Taylor, S. S. Mitosis and apoptosis: how is the balance set? Curr Opin Cell Biol 25, 780–785 (2013).

23. Topham, C. et al. MYC Is a Major Determinant of Mitotic Cell Fate. Cancer Cell 28, 129–140 (2015).

24. Chattopadhyay, S. et al. Role of interferon regulatory factor 3-mediated apoptosis in the establishment and maintenance of persistent infection by Sendai virus. J Virol 87, 16–24 (2013).

25. Mitchison, T. J. The proliferation rate paradox in antimitotic chemotherapy. Mol Biol Cell 23, 1–6 (2012).

26. Weaver, B. A. How Taxol/paclitaxel kills cancer cells. Mol Biol Cell 25, 2677–2681 (2014).

27. Konno, H., Konno, K. & Barber, G. N. Cyclic dinucleotides trigger ULK1 (ATG1) phosphorylation of STING to prevent sustained innate immune signaling. Cell 155, 688–698 (2013).

28. Wheelock, M. S., Wynne, D. J., Tseng, B. S. & Funabiki, H. Dual recognition of chromatin and microtubules by INCENP is important for mitotic progression. J Cell Biol 216, 925–941 (2017).

29. Zeng, X. et al. Pharmacologic inhibition of the anaphase-promoting complex induces a spindle checkpoint-dependent mitotic arrest in the absence of spindle damage. Cancer Cell 18, 382–395 (2010).

30. Gao, P. et al. Structure-function analysis of STING activation by c[G(2',5')pA(3‘,5’)p] and targeting by antiviral DMXAA. Cell 154, 748–762 (2013).

31. Prescott, D. M. & Bender, M. A. Synthesis of RNA and protein during mitosis in mammalian tissue culture cells. Exp Cell Res 26, 260–268 (1962).

32. Bensaude, O. Inhibiting eukaryotic transcription: Which compound to choose? How to evaluate its activity? Transcription 2, 103–108 (2011).

33. Diaz-Martinez, L. A. et al. Genome-wide siRNA screen reveals coupling between mitotic apoptosis and adaptation. EMBO J 33, 1960–1976 (2014).

34. Doménech, E. et al. AMPK and PFKFB3 mediate glycolysis and survival in response to mitophagy during mitotic arrest. Nat Cell Biol 17, 1304–1316 (2015).

35. Orth, J. D., Loewer, A., Lahav, G. & Mitchison, T. J. Prolonged mitotic arrest triggers partial activation of apoptosis, resulting in DNA damage and p53 induction. Mol Biol Cell 23, 567–576 (2012).

36. Orthwein, A. et al. Mitosis inhibits DNA double-strand break repair to guard against telomere fusions. Science 344, 189–193 (2014).

37. Holland, A. J. & Cleveland, D. W. Losing balance: the origin and impact of aneuploidy in cancer. EMBO Rep 13, 501–514 (2012).

38. Garsed, D. W. et al. The architecture and evolution of cancer neochromosomes. Cancer Cell 26, 653–667 (2014).

39. Hayashi, M. T., Cesare, A. J., Rivera, T. & Karlseder, J. Cell death during crisis is mediated by mitotic telomere deprotection. Nature 522, 492–496 (2015).

40. Barber, G. N. STING: infection, inflammation and cancer. Nat. Rev. Immunol. 15, 760–770 (2015).

41. Guse, A., Carroll, C. W., Moree, B., Fuller, C. J. & Straight, A. F. In vitro centromere and kinetochore assembly on defined chromatin templates. Nature 477, 354–358 (2011).

42. Ran, F. A. et al. Genome engineering using the CRISPR-Cas9 system. Nat Protoc 8, 2281–2308 (2013).

43. Held, M. et al. CellCognition: time-resolved phenotype annotation in high-throughput live cell imaging. Nat. Methods 7, 747–754 (2010).

44. Kranzusch, P. J., Lee, A. S.-Y., Berger, J. M. & Doudna, J. A. Structure of human cGAS reveals a conserved family of second-messenger enzymes in innate immunity. Cell Reports 3, 1362–1368 (2013).

45. Lowary, P. T. & Widom, J. New DNA sequence rules for high affinity binding to histone octamer and sequence-directed nucleosome positioning. J Mol Biol 276, 19–42 (1998).

46. Murray, A. W. Cell cycle extracts. Methods Cell Biol 36, 581–605 (1991).

